# Direct anti-inflammatory actions of *N,N*-dimethyltryptamine on microglia are revealed by proteomic profiling and receptor pharmacology

**DOI:** 10.64898/2026.08.05.742931

**Authors:** István Pesti, Ágnes Bessenyei, Rita Frank, Zsuzsanna Darula, Szabolcs Dvorácskó, Zoltán Páhi, Tibor Pankotai, Éva Hunyadi-Gulyás, Krisztián Vinga, Sára Pető, Kata Klein, Ferenc Bari, Ákos Menyhárt, Nicholas V. Cozzi, Eszter Farkas

## Abstract

*N,N*-dimethyltryptamine (DMT) is an endogenous psychedelic tryptamine that has recently emerged as a promising therapeutic candidate for acute ischemic stroke. Although DMT consistently reduces infarct size, attenuates neuroinflammation, and improves functional outcome in experimental stroke, the cellular and receptor mechanisms underlying these effects remain poorly understood.

Primary rat microglial cultures were used to examine the direct anti-inflammatory effects of DMT following lipopolysaccharide (LPS)-induced activation. Microglial morphology, phagocytosis, and proteomic alterations were analyzed. Radioligand binding assays determined the affinity of DMT for microglial sigma-1 receptors (Sig-1Rs). Pharmacological inhibition of Sig-1Rs and serotonin (5-HT) receptors was performed to define receptor-specific mechanisms. Translational relevance was evaluated in acute mouse brain slices subjected to mild oxygen-glucose deprivation (mOGD) and anoxic episodes, where microglial activation, spreading depolarizations (SDs), and neuronal injury were assessed.

DMT directly suppressed LPS-induced microglial activation, promoted a homeostatic morphology, and reduced phagocytic activity. Proteomic profiling demonstrated that DMT selectively reprogrammed inflammatory pathways by suppressing proteins involved in cytokine and chemokine signaling and oxidative stress while largely preserving arachidonic acid–prostaglandin synthesis. DMT bound microglial Sig-1Rs with micromolar affinity comparable to that reported in whole-brain preparations. Pharmacological inhibition revealed that DMT-induced morphological reprogramming required both Sig-1R and serotonergic signaling, whereas suppression of phagocytosis was largely independent of either receptor pathway. In acute brain slices, DMT attenuated microglial activation, reduced SD propagation and ischemic neuronal injury, and tissue-level neuroprotection depended on serotonergic signaling.

DMT directly targets microglia and selectively remodels inflammatory states rather than broadly suppressing microglial activation. The receptor mechanisms underlying its actions are context dependent, with Sig-1R and serotonergic signaling contributing differentially according to the cellular response and experimental model. These findings provide mechanistic insight into the neuroprotective actions of DMT and support its ongoing clinical translation as a potential therapy for ischemic stroke.

## INTRODUCTION

Interest in the therapeutic application of *N,N*-dimethyltryptamine (DMT), the psychedelic compound responsible for the hallucinogenic effects of the traditional South American ceremonial brew ayahuasca (Schultes et al., 1992), has recently gained momentum as a potential therapy for acute ischemic stroke. Preclinical studies have demonstrated that DMT reduces infarct size (Nardai et al., 2020), suppresses spreading depolarizations (SDs) (Szabó et al., 2021), attenuates neuroinflammation and blood-brain barrier disruption (Nardai et al., 2020; László et al., 2025), and improves motor functional outcome (Nardai et al., 2020). These findings have led to the initiation of a Phase I clinical trial (van der Heijden et al., 2025) and the design of a Phase II trial evaluating DMT as a therapy to promote post-stroke recovery. Interest in DMT is further fueled by its endogenous presence throughout the animal kingdom, including humans, and by its proposed role as a neurotransmitter (Schimmelpfennig and Jankowiak-Siuda, 2025). Despite these encouraging preclinical findings and the rapid clinical translation, the mechanisms underlying DMT’s beneficial effects remain incompletely understood.

Many of the neuroprotective effects of DMT have traditionally been attributed to activation of intracellular sigma-1 receptors (Sig-1Rs), which are enriched at the mitochondria-associated endoplasmic reticulum membrane (MAM), a specialized domain where the endoplasmic reticulum is closely apposed to the outer mitochondrial membrane (Frecska et al., 2025). DMT binds Sig-1Rs with low-micromolar affinity (Ki ≈ 14-15 μM) in guinea pig liver and rat brain preparations (Fontanilla et al., 2009; Szabó et al., 2021), and this interaction has long been regarded as a principal mechanism underlying its neuroprotective actions in ischemic brain injury (Nardai et al., 2020; Szabó et al., 2021; Frecska et al., 2025). However, DMT shares its indole ring with serotonin (5-HT) and is also a ligand of several 5-HT receptor subtypes (Schimmelpfennig and Jankowiak-Siuda, 2025). Among these, activation of 5-HT_2A_ receptors is essential for its ability to promote synaptogenesis and neuronal plasticity (Schimmelpfennig and Jankowiak-Siuda, 2025). Notably, 5-HT_2A_ receptor signaling has also been implicated in the modulation of neuroinflammation by DMT (de Deus et al., 2025). Despite these advances, the cellular targets through which DMT exerts its neuroprotective and anti-inflammatory effects within the injured brain remain poorly defined.

Microglia are central regulators of the inflammatory response to cerebral ischemia (Cipriani et al., 2024), making them attractive candidates to mediate the anti-inflammatory effects of DMT. However, current evidence regarding a direct action of DMT on microglia remains inconclusive. While DMT attenuated microglial activation in a recent experimental stroke study (László et al., 2025), an earlier study on global cerebral ischemia found no direct effect on microglia (Szabó et al., 2021). Consequently, whether DMT directly modulates microglial activation remains unresolved, and the accompanying changes in the microglial proteome have yet to be characterized. Moreover, although microglia abundantly express Sig-1Rs (Szabó et al., 2021), the affinity of DMT for microglial Sig-1Rs has not been determined. In addition, microglia express 5-HT receptors (Krabbe et al., 2012), raising the possibility that both Sig-1R- and serotonergic signaling contribute to the potential microglial actions of DMT.

Here, we provide a comprehensive mechanistic analysis of the direct anti-inflammatory actions of DMT on microglia. We first demonstrate that DMT directly suppresses microglial activation in primary cell cultures. Proteomic profiling then identifies molecular pathways underlying the anti-inflammatory and antioxidant actions of DMT by distinguishing protein changes associated with microglial activation that are either sensitive or insensitive to DMT treatment. We next determine the affinity of DMT for microglial Sig-1Rs using radioligand binding assays and combine pharmacological antagonism of Sig-1Rs and 5-HT receptors to define the receptor-specific contributions to distinct aspects of microglial activation. Finally, we show that these direct anti-inflammatory actions extend to an *ex vivo* model of acute ischemia, linking mechanistic observations in cultured microglia with a more physiologically relevant tissue model and strengthening the translational relevance of our findings. Together, our observations establish microglia as a direct cellular target of DMT and provide mechanistic insight into its anti-inflammatory actions in ischemic brain injury.

## MATERIALS AND METHODS

### Ethics declaration

All applicable international, national, and institutional guidelines for the care and use of laboratory animals were followed. All experimental procedures were conducted in accordance with the Hungarian legislation on animal protection (Government Decree 40/2013 (II. 14.) and Act XXVIII of 1998) and the European Union Directive 2010/63/EU on the protection of animals used for scientific purposes. Animals were housed under controlled environmental conditions (20–24 °C, 12-h light/dark cycle) with *ad libitum* access to food and water. Adult mice were deeply anesthetized with 5% isoflurane (N₂O:O₂, 2:1) before decapitation and brain removal, in accordance with the recommendations of the European Commission. No additional ethical approval was required for the *post mortem* collection of brain tissue.

### Drugs

DMT hemifumarate was synthesized at the University of Wisconsin School of Medicine and Public Health, Madison, WI, USA under a State of Wisconsin Controlled Substances Board Special Use Authorization (2130-454) and a Federal Drug Enforcement Administration Schedule 1 license (RC0540751) as previously described (Cozzi and Daley, 2020). Analytical characterization showed that the drug was minimally 99.9% pure.

### Primary microglia cell cultures

#### Maintenance and treatment

Primary cortical microglia co-cultures were prepared from neonatal rats, and microglial monocultures were subsequently isolated as previously described (Pesti et al., 2025). Briefly, cerebral cortices from male and female postnatal day 1 (P1) Sprague-Dawley rats were rapidly dissected, pooled, minced, and dissociated in 0.25% trypsin (Gibco, Thermo Fisher Scientific, Waltham, MA, USA) for 10 min at 37 °C. Enzymatic digestion was terminated by adding Dulbecco’s modified Eagle’s medium (DMEM; Gibco, Thermo Fisher Scientific, Waltham, MA, USA) containing 1 g/L D-glucose, sodium pyruvate, GlutaMA°™, phenol red, 100 U/mL penicillin, 100 μg/mL streptomycin, 0.25 μg/mL amphotericin B, and 10% heat-inactivated fetal bovine serum (FBS; Capricorn Scientific, Ebsdorfergrund-Dreihausen, Germany). Cells were pelleted by centrifugation (1000 × g, 10 min, room temperature), resuspended in DMEM supplemented with 10% FBS, centrifuged again under the same conditions, and filtered through a sterile 100-μm cell strainer (Greiner Bio-One, Kremsmünster, Austria) to remove undissociated tissue fragments. The resulting cell suspension was seeded onto poly-D-lysine-coated T75 culture flasks (1 × 10⁷ cells/flask; Greiner Bio-One) and maintained at 37°C in a humidified incubator with 5% CO₂. The culture medium was replaced the following day and every 3 days thereafter. After 7 days *in vitro* (DIV7), microglia were detached from the mixed glial cultures by orbital shaking (150 rpm, 30 min, 37 °C). The collected supernatant was centrifuged (3000 × g, 8 min, room temperature), and the cell pellet was resuspended in DMEM containing 10% FBS. Cell numbers were determined using a Bürker counting chamber. Microglia were then seeded onto poly-L-lysine-coated coverslips (15 × 15 mm; 2 × 10⁵ cells/coverslip) for immunocytochemistry. The culture medium was replaced the following day and again on days 3 and 6 after subcloning (subDIV6).

#### Activation and pharmacological treatment protocols

To determine the effects of DMT on microglial activation and identify its lowest effective concentration, primary microglial monocultures (subDIV6) were challenged with lipopolysaccharide (LPS; dissolved in DMEM; final concentration: 20 ng/mL; Sigma, St. Louis, MO, USA) (Kata et al., 2016) and treated on day 6 with DMT hemifumarate (dissolved in saline; final concentrations: 5, 10, 20, or 50 μM). The following experimental groups were established: (i) control (unchallenged, untreated), (ii) LPS (challenged with LPS alone), and (iii) LPS+DMT (5, 10, 20, or 50 μM). All treatments were applied for 24 h.

To investigate the receptor specificity of DMT, primary cortical co-cultures (DIV6) and primary microglial monocultures (subDIV6) were used. LPS-challenged cultures were treated with DMT (20 μM) alone or in combination with either the selective Sig-1R antagonist NE-100 (dissolved in distilled water; final concentration: 10 μM; MedChemExpress, USA) or the non-selective serotonin receptor antagonist asenapine (dissolved in distilled water; final concentration: 10 μM; Sigma-Aldrich, USA). The following experimental groups were established: (i) control (unchallenged, untreated), (ii) LPS, (iii) LPS + DMT (20 μM), (iv) LPS + DMT + NE-100, and (v) LPS + DMT + asenapine. All treatments were applied for 24 h.

#### Iba1 immunocytochemistry

Immunocytochemistry was performed as previously described (Kata et al., 2016; Pesti et al., 2025). Briefly, primary cortical co-cultures (DIV7) and microglial monocultures (subDIV7) grown on poly-L-lysine-coated coverslips were fixed with 4% formaldehyde in 0.05 M phosphate-buffered saline (PBS; pH 7.4) for 5 min, washed in PBS (3 × 5 min), and permeabilized and blocked in PBS containing 5% normal goat serum (Sigma-Aldrich, USA) and 0.3% Triton X-100 for 60 min at 37 °C. Cells were then incubated overnight at 4 °C with rabbit anti-Iba1 polyclonal antibody (Abcam, Cambridge, UK; 1:1000) diluted in PBS containing 1% bovine serum albumin (BSA; Sigma-Aldrich, USA) and 0.3% Triton X-100. Following three PBS washes (5 min each), cultures were incubated for 2 h at room temperature in the dark with Alexa Fluor™ 568-conjugated goat anti-rabbit IgG (Invitrogen, Carlsbad, CA, USA; 1:1000) diluted in the same solution without Triton X-100. After three additional PBS washes and a final rinse in distilled water, coverslips were air-dried and mounted with Fluoromount-G containing DAPI (Thermo Fisher Scientific, USA). Images were acquired using a Leica DFC250 camera mounted on a Leica DM LB2 fluorescence microscope (Leica Microsystems, Wetzlar, Germany) equipped with a 40× objective.

#### In vitro phagocytosis assay

The fluid-phase phagocytic activity of microglia in primary cortical co-cultures (DIV7) and microglial monocultures (subDIV7) was assessed by uptake of fluorescent microspheres (2 μm diameter; Sigma, St. Louis, MO, USA). Briefly, fluorescent microspheres were added to the cultures at a final concentration of 0.5 μL/mL from a 2.5% aqueous suspension, and the cultures were incubated at 37 °C for 60 min. Cells were then washed with 2 mL PBS to remove non-internalized microspheres and fixed with 4% formaldehyde in 0.05 M PBS (pH 7.4, room temperature). Samples were subsequently immunolabeled for Iba1 and counterstained with DAPI, as described above (Szabo and Gulya, 2013; Pesti et al., 2025).

#### Image analysis

Digital images of Iba1-immunolabeled preparations were acquired using a Leica DFC250 camera (Leica Microsystems Wetzlar GmbH, Wetzlar, Germany) mounted on a fluorescence microscope (Leica DM LB2; Leica Microsystems CMS GmbH, Wetzlar, Germany) and controlled with LAS X software (Leica Microsystems CMS GmbH, Wetzlar, Germany). Approximately 20 randomly selected cells per experimental condition were analyzed in each of three biological replicates to determine the lowest effective DMT concentration.

Microglial cell silhouettes were generated by converting fluorescence microscopy images of Iba1-immunoreactive cells into binary images using Adobe Photoshop CS3 (Adobe Systems Inc., San Jose, CA, USA). Following binary conversion, the perimeter and area of individual cells were measured using ImageJ (National Institutes of Health, Bethesda, MD, USA). The transformation index (TI), reflecting the degree of process extension, was calculated as perimeter²/(4π × area), as previously described (Pesti et al., 2025). A total of 189 cell silhouettes were analyzed in the receptor antagonist study.

To assess phagocytic activity, cultures were incubated with fluorescent microbeads, and microglia were subsequently identified by Iba1 immunocytochemistry. The proportion of phagocytic microglia and the number of internalized microbeads per phagocytic cell were determined. Twenty non-overlapping random fields were captured from each culture using a Leica DMLB epifluorescence microscope equipped with a 20× objective, and the microbead load of 100–140 cells per culture was quantified using the ImageJ Cell Counter plugin (National Institutes of Health, Bethesda, MD, USA).

#### LC-MS/MS proteomics

Three biological replicates were used in each experimental group (i.e. control, LPS and LPS+20 µM DMT). Primary microglial monoculture samples were digested with trypsin according to the Strap micro protocol (https://files.protifi.com/protocols/s-trap-micro-long-4-7.pdf, accessed on 23 August 2023). Briefly, samples were supplemented with 20% SDS and dried down then redissolved in 50 mM TEAB to give a final SDS concentration of 5% followed by reduction using TCEP (Tris(2-carboxyethyl)phosphine) for 15 min at 37 °C, and alkylation with MMTS (*S*-methyl methanethiosulfonate) for 15 min at room temperature before digesting with MS-grade trypsin (Pierce Biotechnology, Rockford, IL, USA) for 2 h at 47 °C. 10% of the resulting peptide mixtures were loaded onto C18 EvoTips (Evosep) for LC-MS analysis. Reversed-phase separation of the peptides was performed using an Evosep One HPLC (Evosep) applying the „15 SPD” 88-min gradient using a “performance” column packed with 1.5 µm particles of ReproSil-Pur C18 beads (length: 150 mm, internal diameter: 0.15 mm), followed by data-dependent MS/MS acquisition using an Orbitrap Fusion Lumos Tribrid (Thermo Scientific) mass spectrometer equipped with a FAIMS Pro ion mobility device (Thermo Scientific). Data were collected using two compensation voltages (−70 and −50 V) in alternating 1.5 s cycles. All data were acquired with high resolution (120000 and 15000 for MS and MS/MS data, respectively) in the Orbitrap analyzer. AGC target was set at 400,000 and 50,000 for MS and MS/MS acquisition, respectively; MS2 intensity threshold was set to 50,000, higher-energy collision dissociation (HCD) activation was applied using stepped collision energies of NCE = 32 and 35. Dynamic exclusion was enabled (exclusion time: 60 s).

#### In vitro radioligand binding assay

Primary microglial monocultures were homogenized in 20 volumes of ice-cold 50 mM Tris-HCl buffer (pH 8.0 at 25 °C) using a Braun Teflon-glass homogenizer at maximum speed for 30 s. The homogenate was centrifuged at 48,000 × g for 10 min at 4 °C. The resulting pellet was resuspended in 50 mM Tris-HCl buffer (pH 8.0) and centrifuged again. This washing procedure was repeated twice. The final membrane pellet was resuspended in 20 volumes of 50 mM Tris-HCl buffer (pH 8.0), aliquoted, and stored at −80 °C. Protein concentration was determined using the Bradford assay, and samples were diluted to the appropriate protein concentration for the binding assay.

Sig-1R binding assays were performed according to previously established methods with slight modifications (Ishima et al., 2014). Samples were incubated at 27 °C for 120 min in 50 mM Tris-HCl binding buffer (pH 8.0) in a total assay volume of 1 mL containing 0.5 mg/mL membrane protein. Competition binding experiments were performed by incubating microglial membrane preparations with 3.5 nM [³H]-pentazocine in the presence of increasing concentrations (10⁻¹³–10⁻¹¹ M) of unlabeled competing ligands. Non-specific binding was determined in the presence of 10 μM haloperidol. Ki values were calculated using the Cheng–Prusoff equation, applying the previously reported Kd value (13.1 nM) for [³H]-pentazocine (Ishima et al., 2014; Szabó et al., 2021, Kecskés et al. 2025). Binding reactions were terminated by dilution with ice-cold wash buffer (50 mM Tris-HCl, pH 8.0), followed by rapid filtration through 0.1% polyethyleneimine-pretreated Whatman GF/B glass fiber filters (Whatman Ltd., Maidstone, UK) using a 24-well Brandel cell harvester (Brandel, Gaithersburg, MD, USA). Filters were air-dried, immersed in Ultima Gold MV scintillation cocktail, and radioactivity was measured using a TRI-CARB 2100 TR liquid scintillation analyzer (Packard, PerkinElmer, Waltham, MA, USA).

### Acute brain slice preparations

#### Mild oxygen-glucose deprivation and pharmacological treatments

Coronal brain slices were prepared as previously described (Frank et al., 2021). Briefly, adult C57BL/6 mice (n = 18) of both sexes were deeply anesthetized with 5% isoflurane (N₂O:O₂, 2:1) and decapitated. Coronal brain slices (350 μm thick; n=71), corresponding to the striatal region anterior to bregma, were cut using a vibrating blade microtome (Leica VT1000S, Leica, Germany) and collected in ice-cold, modified artificial cerebrospinal fluid (aCSF) containing (in mM): 130 NaCl, 3.5 KCl, 1 NaH₂PO₄, 24 NaHCO₃, 1 CaCl₂, 3 MgSO₄, and 10 D-glucose. Three to five slices were allowed to recover in carbogenated (95% O₂/5% CO₂) normal aCSF containing (in mM): 130 NaCl, 3.5 KCl, 1 NaH₂PO₄, 24 NaHCO₃, 3 CaCl₂, 1.5 MgSO₄, and 10 D-glucose. Subsequently, randomly selected slices were transferred to an interface-type recording chamber (Brain Slice Chamber BSC1, Scientific Systems Design Inc., Ontario, Canada) and continuously superfused with carbogenated aCSF at a flow rate of 2.5 mL/min. The chamber temperature was maintained at 32 °C using a proportional temperature controller (PTC03, Scientific Systems Design Inc., Ontario, Canada).

Brain slices were pre-incubated with the pharmacological agents for 30 min, while slices superfused with normal aCSF served as controls. The pharmacological agents remained present throughout the subsequent experimental protocol. All slices were then subjected to mild oxygen-glucose deprivation (mOGD) by superfusion with aCSF containing 5 mM D-glucose (50% of the normal glucose concentration), as previously described (Frank et al., 2024). After a further 10 min, two SDs were induced by transient withdrawal of oxygen from the gas mixture for 3 min, with an inter-SD interval of 15 min. Recordings were then terminated, and the brain slices were processed for immunocytochemical staining. Similar to the cell culture experiments, the following experimental groups were established to investigate the effects of DMT on the measured variables and to explore the underlying mechanisms using different pharmacological approaches: (i) mOGD (n = 25), (ii) mOGD+DMT (n = 16), (iii) mOGD+DMT+NE-100 (n = 12), and (iv) mOGD+DMT+asenapine (n = 9). All pharmacological agents were dissolved in aCSF to achieve the following final concentrations: DMT, 50 μM; NE-100, 10 μM; and asenapine, 1 μM.

#### Intrinsic optical signal imaging

For intrinsic optical signal (IOS) imaging, slices were illuminated by a halogen lamp (Volpi AG, Intralux 5100, Schlieren, Switzerland). Image sequences were recorded at 1 Hz using a monochrome CCD camera (spatial resolution: 1024×1024 pixel, Pantera 1M30, DALSA, Gröbenzell, Germany) attached to a stereomicroscope (MZ12.5, Leica Microsystems, Wetzlar, Germany), providing 6-10× magnification.

#### TTC staining

To determine cell death and tissue viability in brain slices exposed to mOGD, as well as in naïve slices maintained in normal aCSF without exposure to mOGD or pharmacological treatments (n = 52), 2,3,5-triphenyltetrazolium chloride (TTC; Sigma-Aldrich, USA) staining was performed as previously described (Frank et al., 2024). After completion of the recording protocol, brain slices were immediately transferred from the recording chamber to glass vials and incubated in a 2% TTC solution prepared in 0.1 M PBS for 20 min at 37 °C. The slices were subsequently immersed in 4% paraformaldehyde (PFA) and fixed for 24 h. The stained sections were then mounted on microscope slides and coverslipped with glycerol. Images of the cortex were acquired using a Nikon DS-Fi3 camera mounted on a Leica DM2000 LED light microscope (Leica Microsystems GmbH, Germany) with a 20× objective.

#### Iba1 and NeuN immunofluorescence

To evaluate microglial morphology and neuronal density, immunofluorescent labeling for Iba1 and NeuN was performed. A subset of TTC-stained brain slices (350 μm, n = 18) was paraffin-embedded, sectioned at 3 μm using a rotary microtome (Leica RM2235, Leica, Germany), and mounted onto microscope slides. Following deparaffinization and rehydration, sections were blocked with 10% normal goat serum (Sigma-Aldrich, USA) for 1 h at room temperature and incubated overnight at 4 °C with one of the following primary antibodies: rabbit anti-Iba1 (Abcam, Cambridge, UK; 1:1000) or rabbit anti-NeuN (Abcam, ab177487; 1:500). After three washes in PBS, sections were incubated for 2 h at room temperature in the dark with the corresponding secondary antibodies: Alexa Fluor™ 568-conjugated goat anti-rabbit IgG (Invitrogen, Carlsbad, CA, USA; 1:1000) for Iba1 or Alexa Fluor™ 488-conjugated goat anti-rabbit IgG (Thermo Fisher Scientific, A-11034; 1:1000) for NeuN. Following washes in PBS and distilled water, sections were coverslipped with Fluoromount-G containing DAPI (Thermo Fisher Scientific, USA; 00-4959-52). Images were acquired using a Leica DFC250 camera mounted on a Leica DM LB2 fluorescence microscope (Leica Microsystems, Wetzlar, Germany) equipped with a 40× objective for Iba1 imaging and a 20× objective for NeuN imaging.

#### Image analysis

IOS image sequences were analyzed offline using Fiji (ImageJ, National Institutes of Health, Bethesda, MD, USA). Measurements were performed on contrast-enhanced images. The SD area was determined by manually delineating the cortical region affected by SD at its maximal extent and normalizing it to the total cortical area. For immunohistochemical analysis, microglial cell silhouettes were generated as described above for cultured microglia, and the area of individual cell silhouettes was measured using ImageJ (National Institutes of Health, Bethesda, MD, USA). NeuN-positive cells were quantified using intensity thresholding followed by particle analysis in Fiji. Both histological analyses focused on the parietal cortical region affected by SD. NeuN immunoreactivity was expressed as the percentage of the cortical area occupied by NeuN-positive cells.

### Statistical analysis

Statistical analyses of microglial morphology, phagocytic activity, immunocytochemistry, and intrinsic optical signal (IOS) imaging data obtained from brain slices were performed using GraphPad Prism 8.0 (GraphPad Software, San Diego, CA, USA) or SigmaPlot 12.5 (Systat Software Inc., San Jose, CA, USA). Data distribution was assessed using the Shapiro–Wilk test. Normally distributed data were analyzed by one-way analysis of variance (ANOVA) followed by the Holm–Šidák *post hoc* test and are presented as mean ± SD. Non-normally distributed data were analyzed using the Kruskal–Wallis test followed by Dunn’s multiple-comparison test and are presented as box plots. Statistical significance was set at P < 0.05.

For proteomics analysis, peptide identification and quantitation were done using the Proteome Discoverer software (Thermo Scientific, v3.0). Peptides and proteins were identified with Sequest HT search engine using the rat (sp_canonical TaxID=10116, v2023-06-28, 8178 sequences) and bovine (sp_canonical TaxID=9913, v2022-12-14, 6034 sequences) subsets of the Swissprot protein database. Trypsin was specified as enzyme allowing up to two missed cleavage sites. Mass accuracy was set at 5 ppm and 0.02 Da for precursor and fragment ions, respectively. Constant modification was methylthio on Cys residues. Acetylation and/or loss of Met on protein *N*-terminus, oxidation of Met and cyclization of peptide *N*-terminal Gln were set as variable modifications allowing up to 4 variable modifications per peptide. Label free quantitation was performed using precursor MS signal intensities. Unique and razor peptides were considered for pairwise ratio-based comparison of the sample groups (acceptance parameters: Sequest HT Xcorr>1, S/N in MS spectra > 5, retention time window: 5 min, minimum replicate feature≥50%). Proteins identified with high confidence showing |log2FoldChange|>1 (adjusted p-value ≤ 0.01 using background-based t-test) were accepted as differently expressed.

Principal component analysis (PCA) and hierarchical clustering were performed to assess sample similarity and identify potential outliers. Differentially expressed proteins (DEPs) were identified for the following comparisons: (i) LPS versus Control, (ii) LPS + DMT versus Control, and (iii) LPS + DMT versus LPS. In each comparison, DEPs were defined as proteins with a |log2FC| change greater than 1 and an adjusted P value < 0.05. To evaluate the effects of the treatments on biologically relevant pathways, representative proteins from selected pathways were examined in greater detail. Differential expressions were assessed based on log2-fold change and adjusted P values across the three comparisons. The heatmap was generated with ComplexHeatmap 2.14.0 and circlize 0.4.16 in R 4.2.1 from row-wise z-scored log2 abundance ratios, clustered by Euclidean distance and complete linkage. Volcano plots were generated in R 4.2.1 (x86_64-w64-mingw32/x64, Windows) using EnhancedVolcano 1.16.0 with ggplot2 4.0.0 and ggrepel 0.9.6 for non-overlapping labels. The protein-level Proteome Discoverer export was read with readxl 1.4.5 and processed with dplyr 1.1.4; result tables were written with openxlsx 4.2.8. Abundance ratios were log2-transformed and plotted against the adjusted p-values reported by Proteome Discoverer, using pCutoff = 0.01 and FCcutoff = 1. The complete proteomic dataset, including all significantly differentially expressed proteins, is provided in the Supporting Information.

## RESULTS

### DMT inhibits activation of cultured microglia

DMT has recently been shown to mitigate microglial activation in a preclinical model of acute ischemic stroke (László et al., 2025). However, it has remained unclear whether DMT acts directly on microglia or exerts its effects indirectly by targeting neurons, thereby secondarily modulating microglial activation. Therefore, we sought to determine whether DMT directly inhibits microglial activation, using a primary microglial monoculture model previously employed to demonstrate the anti-inflammatory effects of calcium channel antagonism in LPS-challenged microglia (Pesti et al., 2025). In addition, we identified the lowest effective concentration of DMT in this experimental model.

The morphological phenotype of microglia is a sensitive indicator of activation, with low TI values reflecting the ameboid morphology induced by LPS challenge (Pesti et al., 2024). DMT dose-dependently increased the LPS-induced reduction in TI, resulting in a progressively more ramified, resting-state morphology (Fig. 1A).

**Figure 1.**
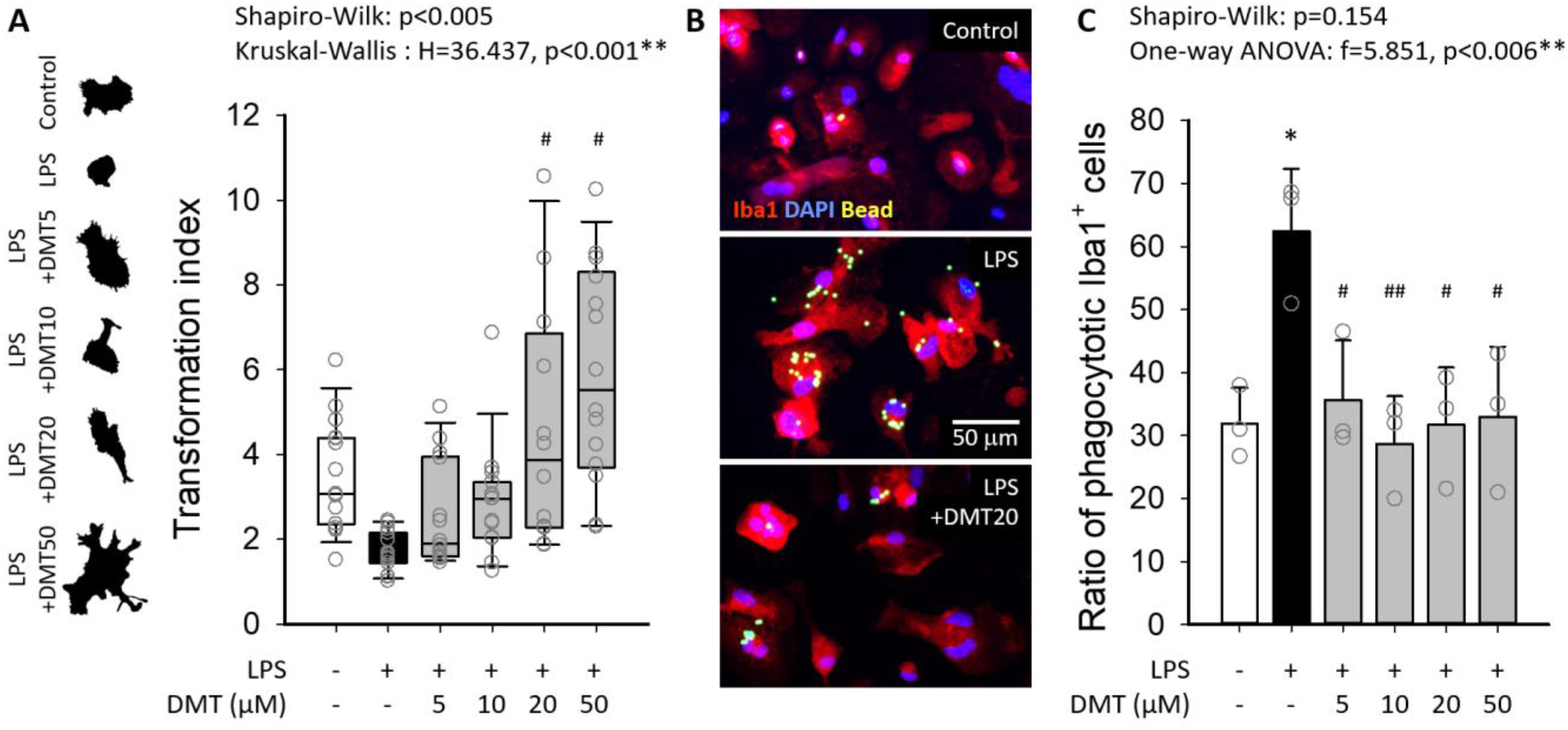
DMT attenuates ameboid transformation and phagocytosis of activated microglia. **(A)** Quantitative analysis of microglial morphology using the transformation index (TI) demonstrates that DMT at concentrations of 20-50 μM reverses the amoeboid transformation induced by LPS. Representative binary silhouettes of microglia for each experimental condition are shown on the left. As the data were not normally distributed, results are presented as box plots with individual data points overlaid. Statistical symbols indicate differences between groups according to Dunn’s post hoc test, with significance levels defined as: p < 0.05* vs. Control, p < 0.05^#^ vs. LPS. (**B)** Representative fluorescence micrographs of Iba1-positive cultured microglia containing phagocytosed fluorescent microbeads. LPS stimulation increases the proportion of phagocytic microglia, whereas DMT inhibits this response. **(C)** LPS nearly doubled the proportion of phagocytic microglia, whereas DMT reduced phagocytosis to control level, with a significant effect observed already at the lowest concentration tested (5 μM). Data are shown as mean ± stdev. Statistical symbols indicate differences between groups according to the Holm-Sidak post hoc test, with significance levels defined as: p < 0.05* vs. Control, and p < 0.05^#^, p < 0.01^##^ vs. LPS.

Compared with LPS treatment alone, the effect of DMT reached statistical significance at 20 μM. The functional phenotype of microglial activation was further assessed by phagocytic activity (Pesti et al., 2024), which was markedly enhanced by LPS. Co-treatment with DMT significantly reduced the proportion of phagocytic microglia, with a significant effect observed already at the lowest concentration tested (5 μM) (Fig. 1B-C). Together, these findings demonstrate that DMT directly promotes the restoration of the resting microglial phenotype following LPS activation. Because the effect of DMT on microglial morphology first reached statistical significance at 20 μM, this concentration was considered the lowest effective dose and selected for subsequent experiments investigating the proteomic changes associated with DMT treatment.

### DMT modulates inflammatory and oxidative stress markers in cultured microglia

In total, 3610 proteins were identified by LC-MS/MS in primary microglial monocultures, of which 3316 were quantified. Across the three pairwise comparisons (LPS vs. control, LPS + DMT vs. control, and LPS + DMT vs. LPS), 134 proteins were differentially expressed (DEPs) (Fig. 2). Activation by LPS upregulated several components of cytokine and chemokine signaling (e.g. interleukin-β (Il1b), log_2_FC = 1.4792*; interleukin 6 signal transducer (Il6st), log_2_FC = 1.0072*; C-C motif chemokine ligand 20 (Ccl20), log_2_FC = 2.0472*) (Fig. 3), notably elevated the expression of proteins associated with nitric oxide production (inducible nitric oxide synthase (Nos2), log_2_FC = 3.3865*, L-arginine transporter (Slc7a2), log_2_FC = 1.2290*), and significantly increased protein levels involved in prostaglandin biosynthesis (secretory phospholipase A2 (Pla2g2a), log_2_FC=1.4895*; cyclooxygenase-2 (Ptgs2), log_2_FC = 2.0824*; prostaglandin E synthase (Ptges), log_2_FC = 1.245*) (Fig. 4). These changes collectively indicate enhanced immune function and linked oxidative stress, as expected.

**Figure 2.**
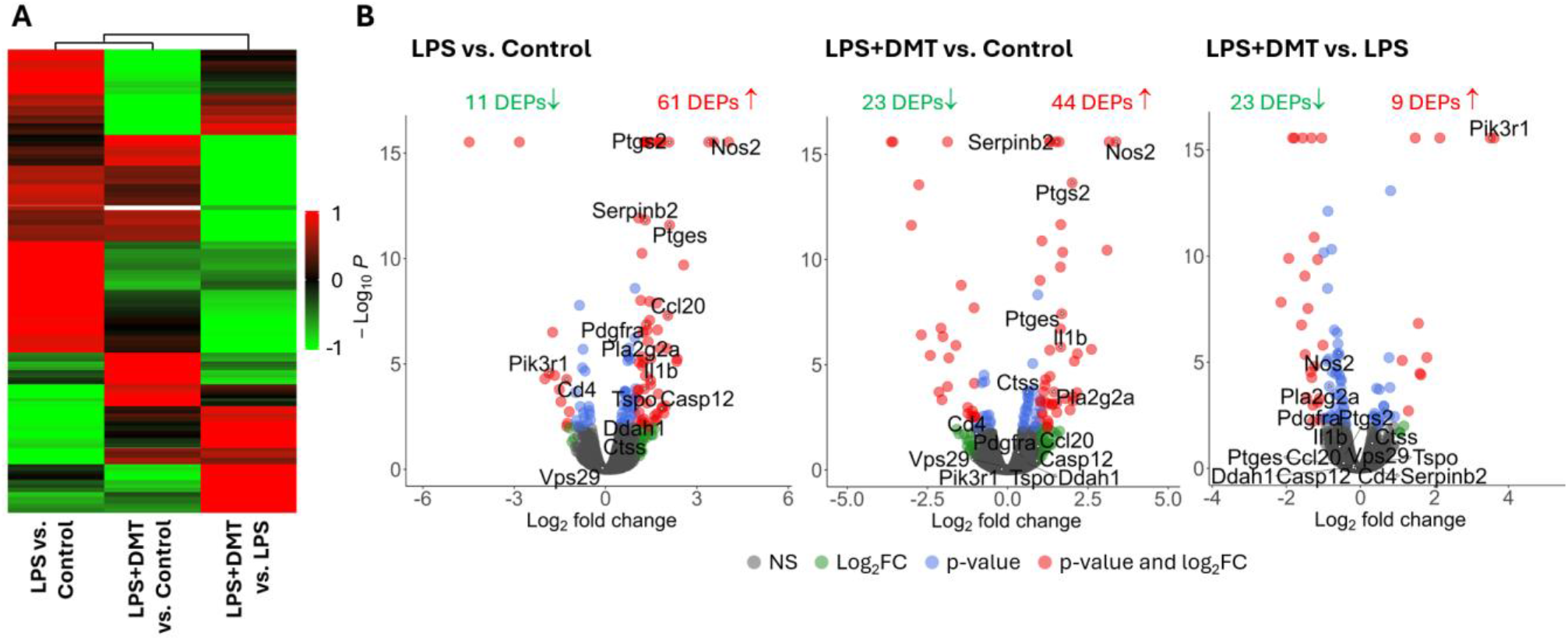
Activation with LPS and treatment with DMT alter the proteome of rat primary microglia. (A) Heat map of all quantified protein groups. Colors represent z-scored log₂ abundance ratios, reflecting the relative pattern of change between comparisons (LPS vs. Control, LPS + DMT vs. Control, LPS + DMT vs. LPS) **(B)**, Volcano plots depicting log₂ fold changes versus –log₁₀ adjusted p-values for each gene in the following comparisons: LPS vs. Control, LPS + DMT vs. Control, and LPS + DMT vs. LPS alone. Cutoff values are |log2FC| = 1 and adjusted p = 0.01.

**Figure 3.**
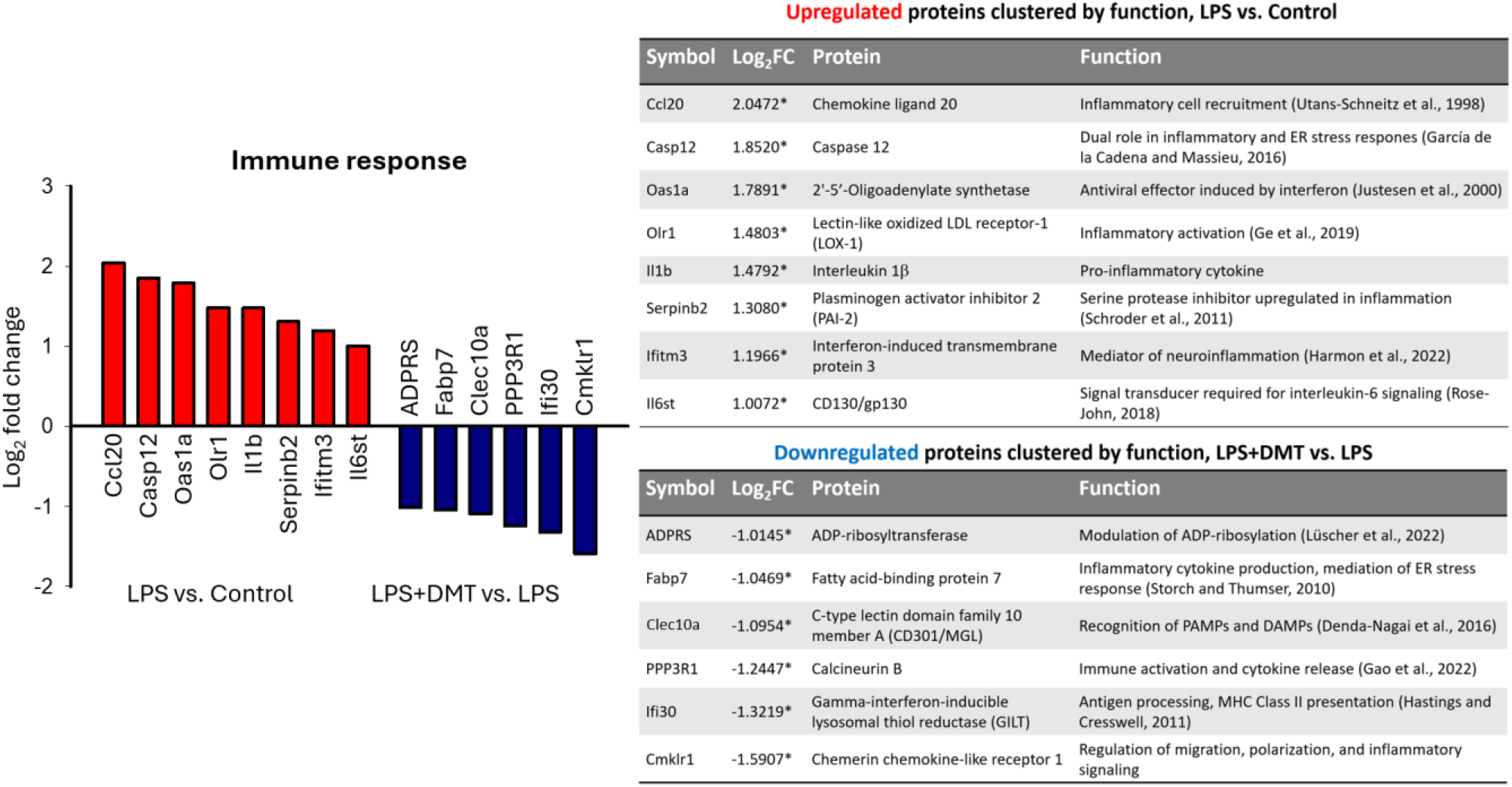
DMT downregulates immune-related proteins that functionally oppose the actions of proteins upregulated by LPS. Protein abundance is represented as log₂ fold changes (Log₂FC), with significance defined as adjusted p < 0.01*. Proteins downregulated by DMT (LPS + DMT vs. LPS; blue) are shown in contrast to those upregulated by LPS (LPS vs. Control; red). Protein symbols are decoded, and their functions are detailed in the accompanying tables.

**Figure 4.**
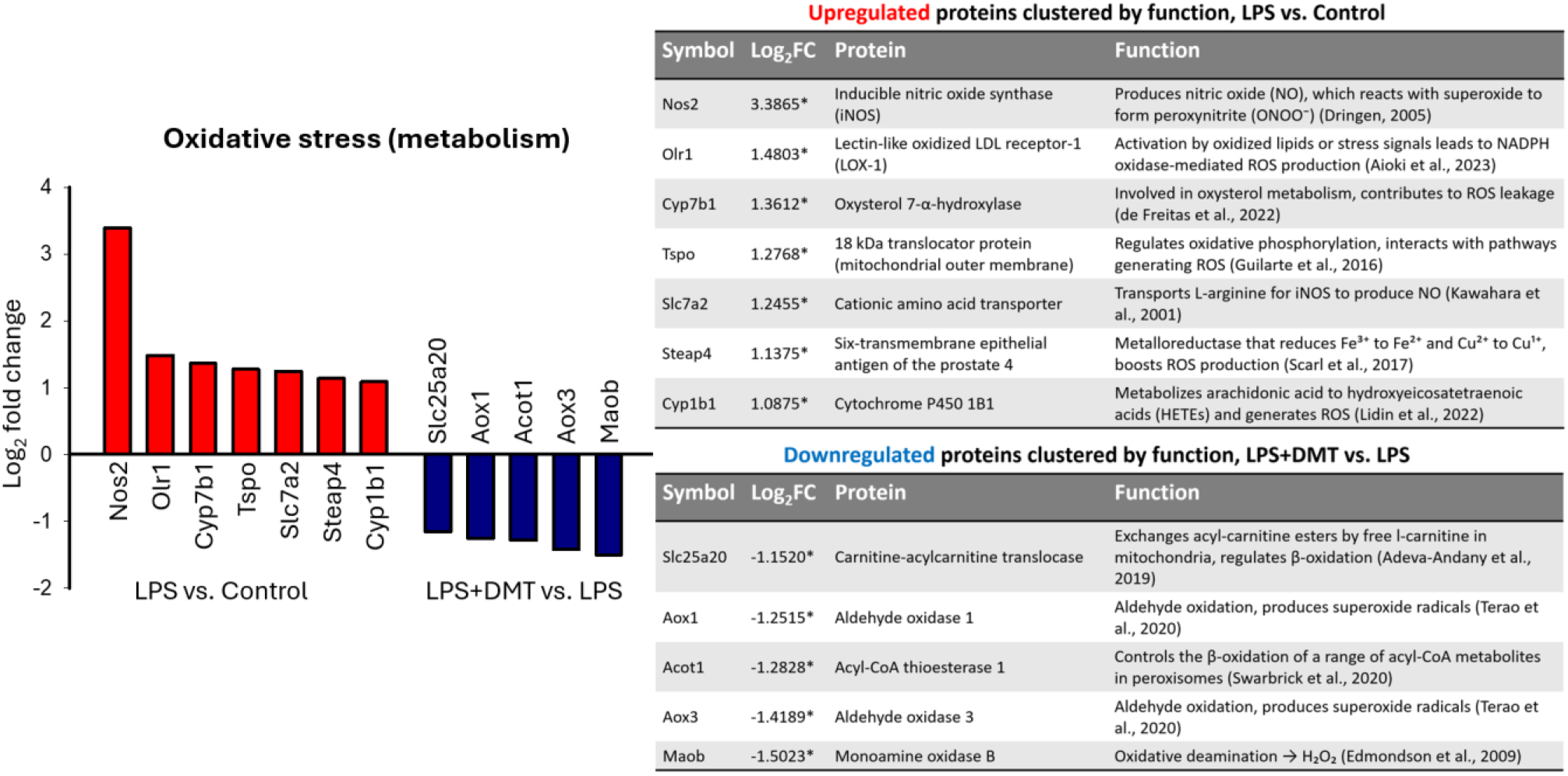
DMT downregulates oxidative stress-related proteins that counterbalance the oxidative stress response induced by LPS. Protein abundance is represented as log₂ fold changes (Log₂FC), with significance defined as adjusted p < 0.01*. Proteins downregulated by DMT (LPS + DMT vs. LPS; blue) are shown in contrast to those upregulated by LPS (LPS vs. Control; red). Protein symbols are decoded, and their functions are detailed in the accompanying tables.

Of the 61 DEPs upregulated by LPS (LPS vs. Control), the increase in abundance of 36, including Il6st and Ccl20, was no longer evident following the addition of DMT (LPS + DMT vs. Control). Conversely, of the 11 DEPs downregulated by LPS (LPS vs. Control), decrease in the levels of 7 was no longer detectable after DMT treatment (LPS + DMT vs. Control). In total, 60% of LPS-responsive DEPs (LPS vs. Control) were no longer detectable upon DMT addition (LPS + DMT vs. Control). Further, DMT appeared to counterbalance processes associated with immune activation and oxidative stress by downregulating protein sets complementary to those activated by LPS (Fig. 3-4). Accordingly, several components of the immune response including proteins involved in the recognition of pathogen- or danger-associated molecular patterns (Clec10a), antigen processing (Ifi30), and inflammatory signaling (Fabp7, PPP3R1, Cmklr1) were suppressed by DMT (LPS + DMT vs. LPS) (Fig. 3). Of note, prostaglandin synthesis pathway proteins (Pla2g2a, Ptgs2, and Ptges), which were upregulated by LPS, remained resistant to DMT, as evidenced by their sustained upregulation following treatment (log_2_FC = 1.4468*, 2.0079* and 1.6857*, respectively; LPS + DMT vs. Control).

Cellular metabolism, which is markedly altered during microglial activation (Sabogal-Guáqueta et al., 2023), is a key source of reactive oxygen species (ROS) during the immune response (Block et al., 2007). In addition to increased enzymatic production of nitric oxide (NO), aldehyde and fatty acid oxidation also contribute to elevated ROS levels. LPS substantially upregulated inducible nitric oxide synthase (iNOS; Nos2) and the cationic amino acid transporter (Slc7a2), which supplies L-arginine for NO synthesis (LPS vs. Control).

The abundance of certain cytochrome P450 family members (Cyp7b1 and Cyp1b1, involved in oxysterol and arachidonic acid metabolism, respectively) was also increased following LPS activation (LPS vs. Control), potentially contributing further to ROS generation. DMT had no effect on the expression of these proteins (except for Cyp1b1), as indicated by their continued upregulation (LPS + DMT vs. Control). However, DMT reduced the abundance of monoamine oxidase B (Maob) and aldehyde oxidases (Aox1 and Aox3), which may represent a compensatory mechanism to limit ROS production via alternative pathways (Fig. 4).

### DMT binds to Sig-1R in cultured microglia

In the context of neuroprotection, DMT has been proposed to exert its beneficial effects through activation of Sig-1Rs (Szabó et al., 2021; Frecska et al., 2025; László et al., 2025). To determine whether DMT binds to Sig-1Rs expressed by microglia, thereby supporting this hypothesis, we performed competition binding assays using rat microglia homogenates and the Sig-1R-specific radioligand [³H]-pentazocine. The experiments revealed that DMT binds to Sig-1R with a Ki value of 3.2 µM (Fig. 5). The well-known Sig-1R ligand (+)-pentazocine showed high binding affinity, with a Ki value of 9.8 nM (Fig. 5). Specific binding accounted for 65% of total binding at a radioligand concentration of 3.5 nM under equilibrium conditions. The observed DMT affinity (Ki = 3.2 µM) is in good agreement with previous studies performed on rat brain homogenates (Szabó et al., 2021), or rat liver homogenates (Fontanilla et al., 2009), supporting the presence of pharmacologically relevant Sig-1R binding sites in cultured primary rat microglia.

**Figure 5.**
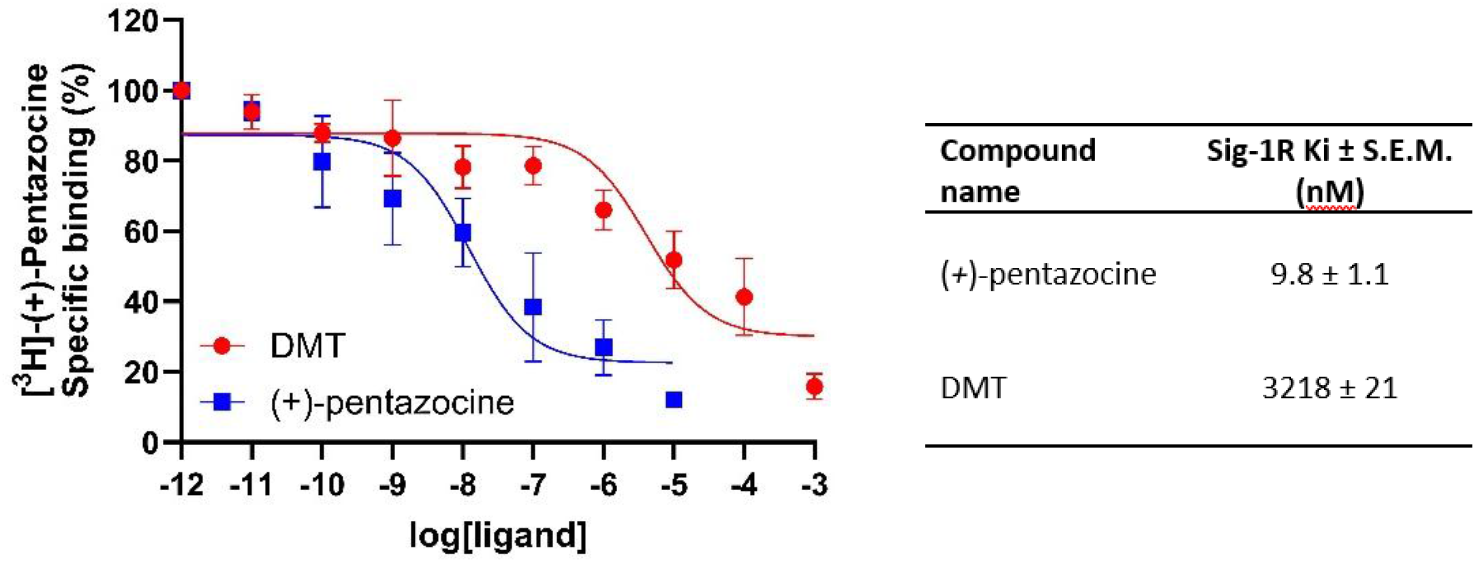
DMT binds to Sig-1R in rat microglia homogenates. Competitive binding curves of DMT against the radioactive Sig-1R ligand [^3^H](+)-pentazocine. Data are expressed as the percentage of specific binding.

### DMT regulates microglial morphology and phagocytosis through distinct receptor mechanisms

To further investigate the receptor targets underlying the effects of DMT, we performed microglial culture experiments in which LPS-activated microglia were treated with DMT alone or in combination with the Sig-1R antagonist NE-100 or the broad-spectrum 5-HT receptor antagonist asenapine. Consistent with our initial experiments (Fig. 1B), DMT at 20 μM attenuated LPS-induced amoeboid transformation in both primary co-cultures and microglial monocultures, as indicated by TI values that did not differ significantly from those of the control cultures (Fig. 6). This effect was more pronounced in co-cultures (Fig. 6A-B). Co-treatment with either NE-100 or asenapine abolished the effects of DMT, reducing TI values to levels characteristic of LPS-activated microglia, particularly in co-cultures (Fig. 6).

**Figure 6.**
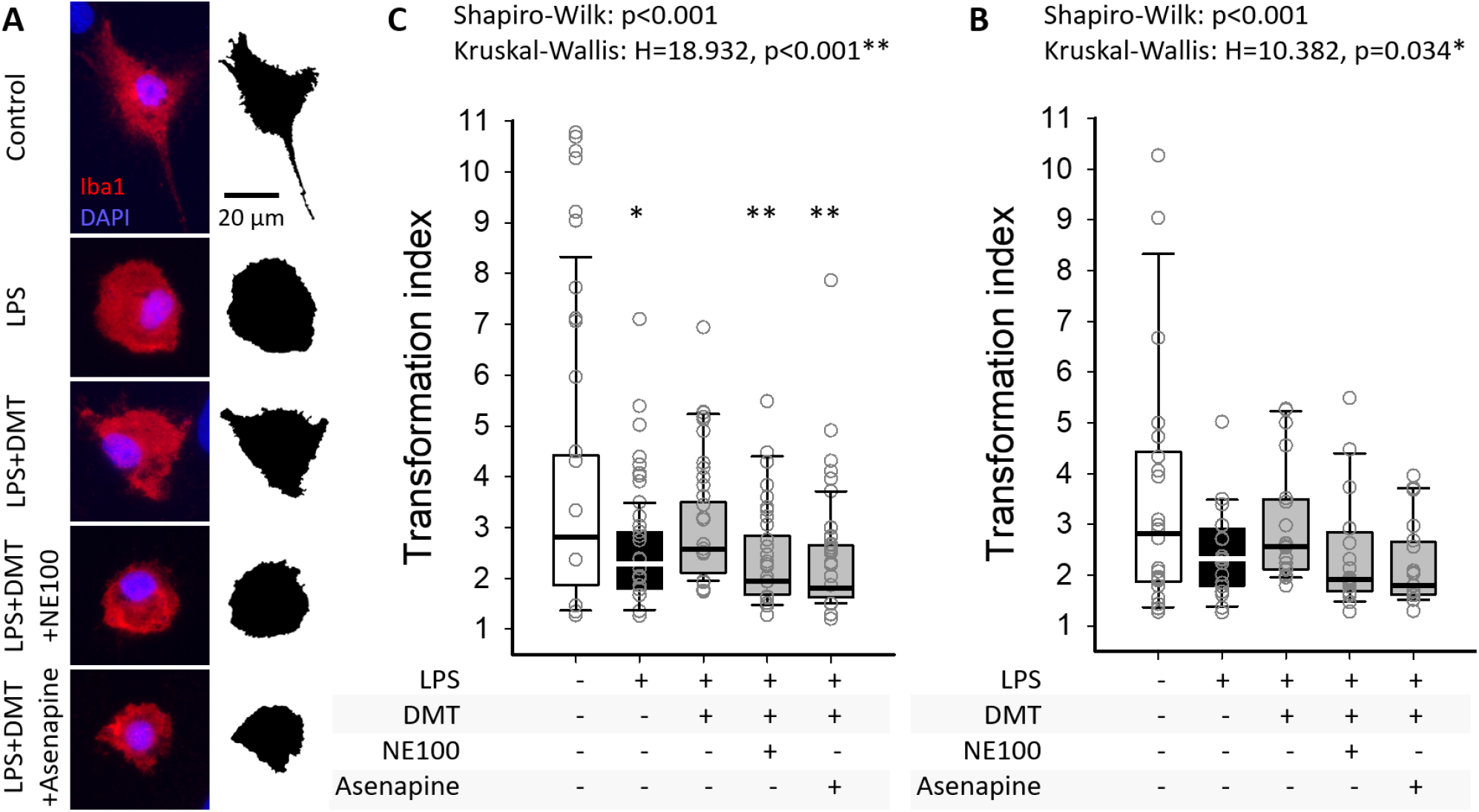
The Sig-1R antagonist NE-100 and the broad-spectrum 5-HT receptor antagonist asenapine reverse the inhibitory effect of DMT on microglial amoeboid transformation. **(A)** Representative fluorescence micrographs of cells from primary co-cultures show Iba1-positive microglia and their corresponding binary silhouettes. LPS induces amoeboid transformation, whereas DMT restores the ramified microglial phenotype. Both NE-100 and asenapine reverse the effects of DMT. **(B)** Quantitative analysis of microglial morphology in primary co-cultures shows that pharmacological blockade of Sig-1R or 5-HT receptors attenuates the effects of DMT. **(C)** Quantitative analysis of microglial morphology in primary microglial monocultures shows similar trends. As the data are not normally distributed, results are presented as box plots, with individual data points overlaid. Statistical symbols indicate differences between groups according to Dunn’s post hoc test, with significance levels defined as: p < 0.05* and p < 0.01** vs. Control.

LPS-induced phagocytosis was reduced by DMT, particularly in microglial monocultures (Fig. 7), consistent with our initial findings (Fig. 1C). Although the proportion of phagocytic microglia increased more markedly in co-cultures than in monocultures following LPS stimulation (from 22% to 66% and from 22% to 34%, respectively), DMT reduced the proportion of phagocytic cells only in monocultures (from 34% to 17%). In contrast to their effects on microglial morphology, neither NE-100 nor asenapine altered this response. Phagocytic activity was further quantified by determining the number of internalized microbeads per microglial cell. Consistent with the changes in the proportion of phagocytic cells, DMT reduced LPS-induced microbead uptake. This effect was not reversed by either receptor antagonist, although asenapine moderately increased phagocytic activity in microglial monocultures relative to DMT alone (Fig. 7).

**Figure 7.**
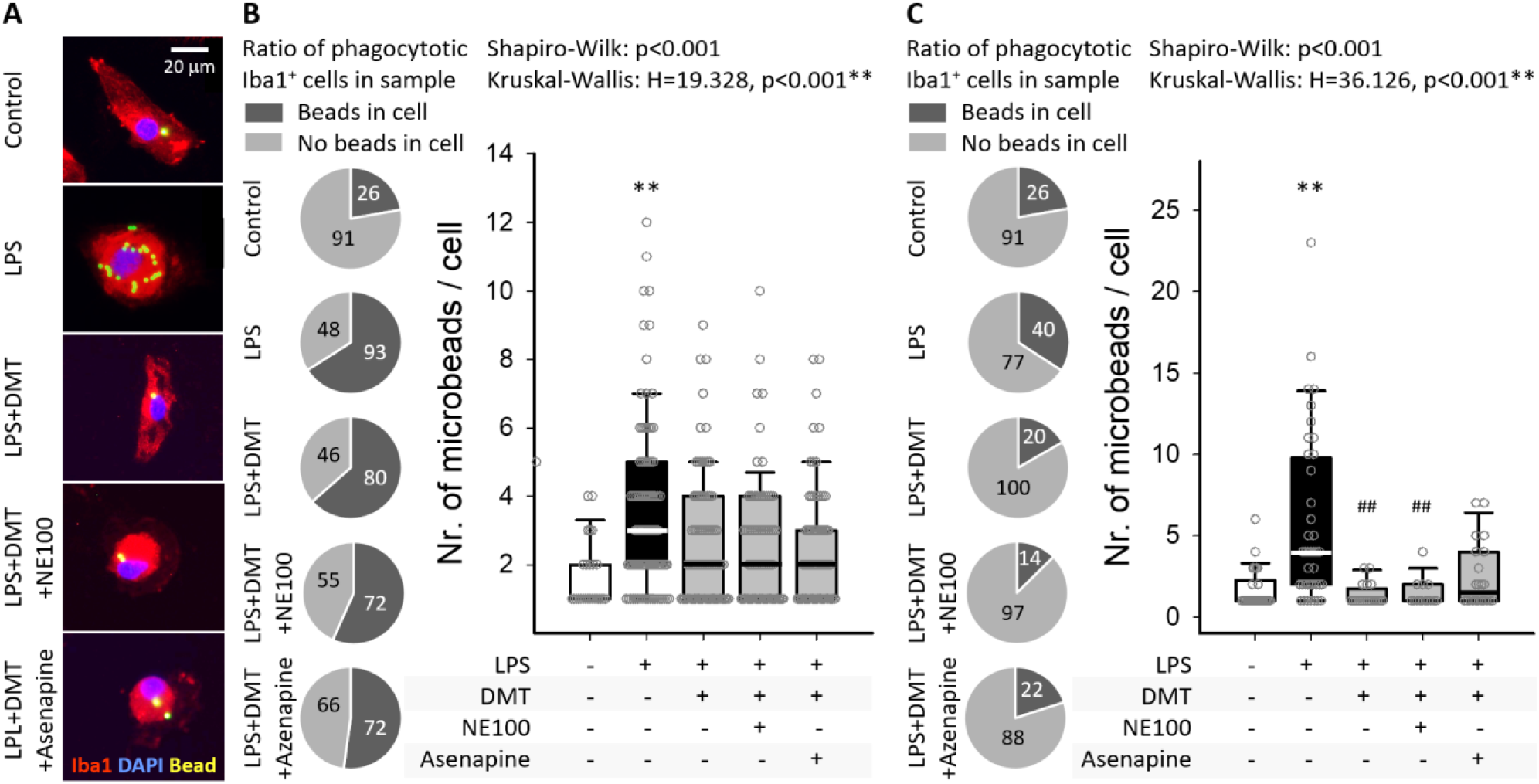
Neither the Sig-1R antagonist NE-100 nor the broad-spectrum 5-HT receptor antagonist asenapine blocks the inhibitory effect of DMT on phagocytosis in activated microglia. **(A)** Representative fluorescence micrographs of primary microglial monocultures show Iba1-positive microglia, containing phagocytosed fluorescent microbeads. LPS stimulation increases the number of internalized microbeads, whereas DMT inhibits this response. Neither NE-100 nor asenapine attenuates the inhibitory effect of DMT on microglial phagocytosis. **(B)** In primary co-cultures, pie charts illustrate the proportion of phagocytic microglia (with number of cells displayed), which is markedly increased by LPS but is not significantly affected by DMT, NE-100, or asenapine. In contrast, quantification of phagocytic activity of individual cells, expressed as the number of internalized microbeads per microglial cell, demonstrates an inhibitory effect of DMT that is not reversed by NE-100 or asenapine. **(C)** In primary microglial monocultures, DMT reduces both the proportion of phagocytic microglia and phagocytic activity, with a pronounced effect. As the quantitative data are discrete numbers and are not normally distributed, results are presented as box plots, with individual data points overlaid. Statistical symbols indicate differences between groups according to Dunn’s post hoc test, with significance levels defined as: p < 0.01** vs. Control and p < 0.01^##^ vs. LPS.

### DMT suppresses SDs, preserves tissue integrity, and attenuates microglial activation in ex vivo brain slices exposed to ischemia

To link the observations obtained in cultured microglia with a more pathophysiologically relevant experimental setting, SD characteristics and histological endpoints were evaluated in live brain slice preparations. Brain slices were challenged by mOGD and two episodes of transient anoxia, the latter serving to trigger SDs in a controlled manner. SDs occurred reliably in response to anoxia, with a latency of 67 ± 42 s for the first SD during anoxia 1 and 33 ± 20 s for the first SD during anoxia 2, indicating increased susceptibility to SD generation during the second anoxic episode. In 26% of the slices during anoxia 1 and 18% during anoxia 2, the initial SD was followed by one or two recurrent SDs (rSDs). Notably, recurrent SDs were not observed during anoxia 2 in DMT-treated slices.

The cortical surface area affected by each SD was expressed relative to the total cortical surface area and summed across all SD events to provide an integrated measure of SD burden, with a smaller cumulative SD area indicating greater inhibition of SD. DMT treatment tended to reduce the cumulative SD area (102 ± 53 vs. 138 ± 52%, DMT vs. mOGD). This effect was unaffected by co-treatment with NE-100 (109 ± 51%) but was abolished by asenapine (136 ± 20%) (Fig. 8A). This trend was further supported by histological analyses. TTC staining, used to assess tissue viability, revealed a marked loss of viable tissue following mOGD, which was partially reversed by DMT. Consistent with the SD findings, co-treatment with NE-100 did not significantly alter the protective effect of DMT, whereas asenapine abolished it (Fig. 8B). Finally, NeuN immunolabeling confirmed the neuroprotective effect of DMT, which was similarly counteracted by co-administration of asenapine (Fig. 8C). Collectively, these findings suggest that tissue injury and SD burden are closely associated, and that the neuroprotective effects of DMT are mediated predominantly through serotonergic signaling.

**Figure 8.**
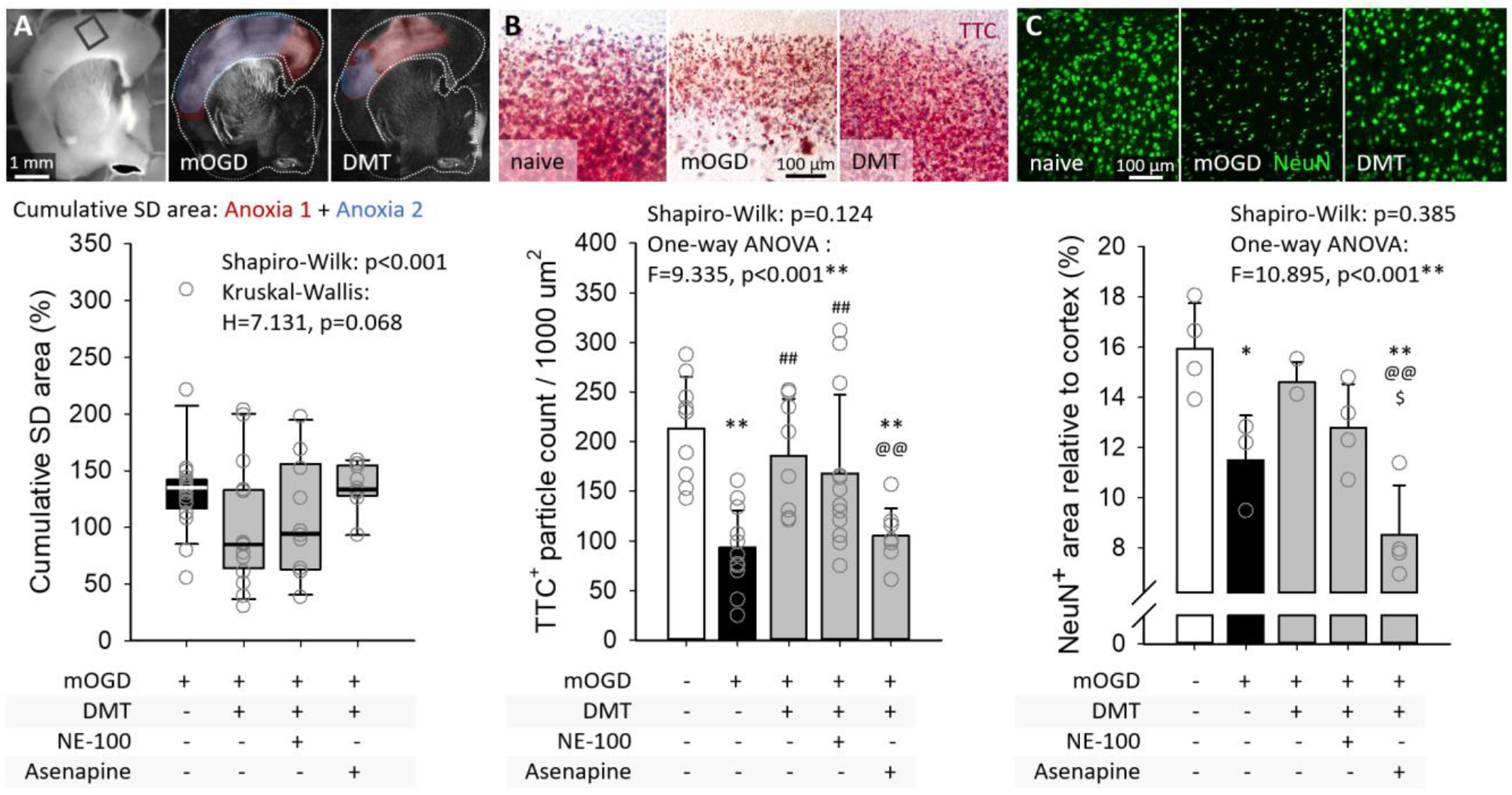
DMT suppresses SD and tissue injury in ischemic/anoxic live brain slice preparations. **(A)** Representative IOS images illustrate the cortical area invaded by SDs during anoxia 1 (red) and anoxia 2 (blue) in the same brain slice exposed to mOGD alone or mOGD with DMT treatment. The rectangle in the first image indicates the region of interest used for the histological analyses shown in panels B and C. The cumulative SD area, calculated as the cortical area affected by each SD relative to the total cortical surface area and summed across both anoxic episodes, showed a trend toward reduction by DMT, which was abolished by asenapine. As the data were not normally distributed, results are presented as box plots with individual data points overlaid. **(B)** Representative TTC-stained brain slices demonstrate tissue injury induced by mOGD and its attenuation by DMT treatment. Quantitative analysis confirmed the protective effect of DMT, which was abolished by co-treatment with asenapine. **(C)** Representative NeuN-immunolabeled sections confirm the TTC findings and demonstrate the neuroprotective effect of DMT. Co-treatment with asenapine again abolished this protective effect. Data in B and C are shown as mean±stdev. Statistical symbols indicate differences between groups according to the Holm-Sidak post hoc test, with significance levels defined as: p < 0.05* and p < 0.01**vs. naive, p < 0.01^##^ vs. mOGD, p < 0.01^@@^ vs. DMT, and p < 0.05^$^ vs. NE-100.

Finally, the effects of DMT on microglial activation were evaluated in the brain slice preparations. Among the morphological parameters examined, the area occupied by individual Iba1-positive microglia, reflecting the degree of process arborization, proved to be the most sensitive indicator of treatment effects. Consistent with the findings obtained in cultured microglia (Fig. 6), DMT promoted a ramified, resting microglial phenotype, as reflected by the increased cell area (1863 ± 247 vs. 1066 ± 109 µm^2^, DMT vs. mOGD) (Fig. 9). As observed in the cell culture experiments, the effects of DMT were abolished by co-treatment with either NE-100 or asenapine (Fig. 9). These findings suggest that both Sig-1R and serotonergic signaling contribute to the inhibitory effects of DMT on microglial activation *in situ*.

**Figure 9.**
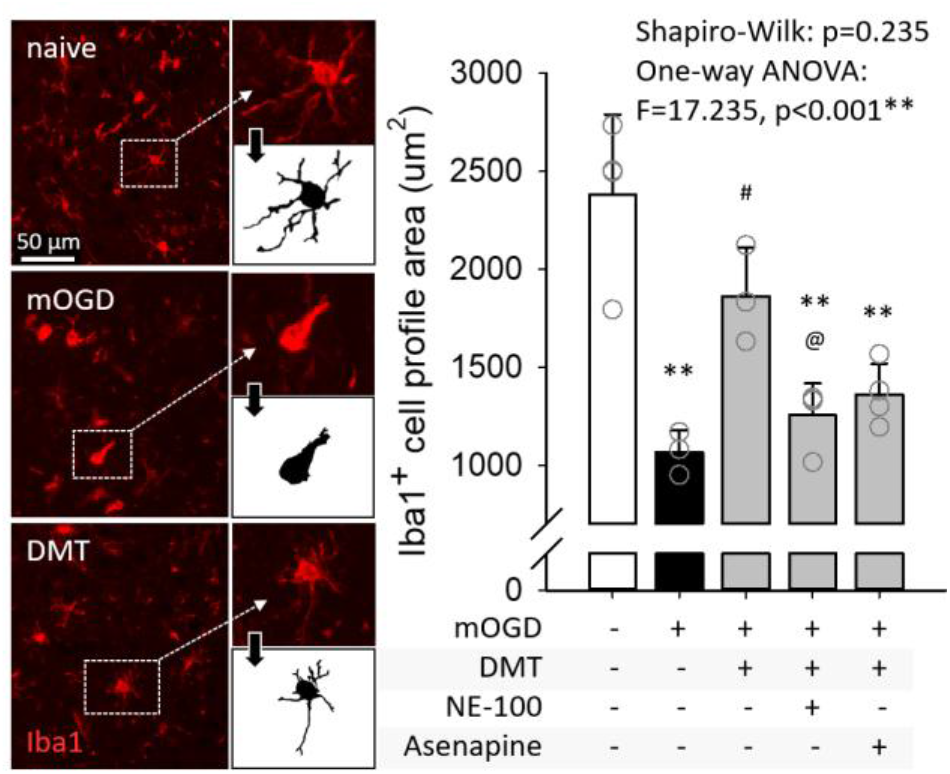
DMT attenuates microglial activation in ischemic/anoxic live brain slice preparations through Sig-1R and 5-HT receptor involvement. Representative Iba1-immunolabeled sections illustrate the morphological phenotype of microglia, with the amoeboid morphology characteristic of mOGD and a more ramified morphology following DMT treatment, indicative of the resting state. Quantitative analysis of the area occupied by Iba1-positive cell profiles demonstrated that DMT increased microglial cell area, whereas co-treatment with either NE-100 or asenapine abolished this effect. Data are shown as mean±stdev. Statistical symbols indicate differences between groups according to the Holm-Sidak post hoc test, with significance levels defined as: p < 0.01**vs. naive, p < 0.05^#^ vs. mOGD, and p < 0.05^@^ vs. DMT.

## DISCUSSION

In the present study, we identify microglia as a direct cellular target of DMT and delineate distinct molecular and receptor mechanisms underlying its anti-inflammatory actions.

LPS-activated microglia in primary cultures, as well as microglia activated by metabolic challenge in live brain slice preparations, underwent a characteristic amoeboid transformation (Fig. 1, Fig. 9). Importantly, DMT treatment reversed this activated phenotype toward a resting morphology in both experimental models and inhibited LPS-induced phagocytosis in primary microglial cultures (Fig. 1, Fig. 9). Together, these findings demonstrate that DMT directly targets microglia and suppresses their activation in various contexts, consistent with the most recent report describing the protective effects of DMT in experimental acute ischemic stroke (László et al., 2025).

Key features of microglial activation include the production of pro-inflammatory cytokines and chemokines, the generation of ROS and oxidative stress, and the synthesis of arachidonic acid-derived lipid mediators. Proteomic analysis revealed that DMT selectively reprograms these inflammatory processes (Fig. 2). Specifically, DMT modulated the expression of proteins involved in inflammatory signaling and oxidative stress (Fig. 3-4) while leaving the expression of key enzymes involved in arachidonic acid–prostaglandin synthesis largely unaffected.

Among the LPS-induced proteins whose expression was reduced by DMT, CCL20 and CMKLR1 may be particularly relevant to ischemic brain injury. Both proteins are upregulated in experimental stroke models (Utans-Schneitz et al., 1998; Terao et al., 2009; Long et al., 2025) and participate in the regulation of leukocyte recruitment, inflammatory cell migration, and intercellular communication (Schutyser et al., 2003; Mariani and Roncucci, 2015). As microglia play a central role in orchestrating leukocyte infiltration into the injured brain, a hallmark of neuroinflammation following ischemic stroke (Planas, 2018), DMT-mediated suppression of these proteins in cultured microglia may therefore have therapeutic relevance in ischemic stroke.

DMT did not attenuate the LPS-induced upregulation of the canonical iNOS/L-arginine pathway, suggesting that its anti-inflammatory effects are not mediated through suppression of nitric oxide synthesis. However, DMT reduced the abundance of monoamine oxidase B (MAOB) and aldehyde oxidases, enzymes that generate ROS during monoamine and aldehyde metabolism. Notably, pharmacological inhibition of MAOB has been shown to exert anti-inflammatory and neuroprotective effects in experimental acute ischemic stroke, at least in part through modulation of microglial activation (Zou et al., 2022). Accordingly, DMT may attenuate oxidative stress by limiting alternative sources of ROS without suppressing the canonical iNOS pathway. Such selective modulation may be advantageous in ischemic stroke, where excessive oxidative stress contributes substantially to secondary tissue injury, whereas nitric oxide signaling is also important for maintaining cerebral vasodilation to promote tissue perfusion (Wierońska et al., 2021).

LPS also upregulated the enzymes of the secretory phospholipase A2–cyclooxygenase-2–prostaglandin E synthase pathway, which mediates the production of the lipid mediator PGE_2_ in activated microglia (Minghetti and Levi, 1995). Interestingly, DMT did not alter the expression of these enzymes. This observation is noteworthy because PGE_2_ exerts context-dependent effects and may even confer neuroprotection in the ischemic brain through activation of PGE_2_ receptors, particularly EP2 and EP4, and subsequent cyclic AMP signaling (McCullough et al., 2004; Ahmad et al., 2006). Taken together, these findings suggest that DMT selectively suppresses inflammatory programs while sparing signaling pathways, such as PGE_2_ synthesis and nitric oxide production, that may retain physiological or even neuroprotective functions during ischemic injury. Since the neuroprotective effects of DMT in ischemic stroke have been proposed to be mediated through Sig-1R activation (Frecska et al., 2025; László et al., 2025), and Sig-1Rs are ubiquitously expressed in microglia (Szabó et al., 2021), we next determined the binding affinity of DMT for Sig-1Rs in cultured microglia, which had not previously been characterized. The measured Ki was within the micromolar range (Fig. 5) previously reported for whole-brain or liver preparations (Fontanilla et al., 2009; Szabó et al., 2021), indicating that microglial Sig-1Rs exhibit a comparable pharmacological profile. Together with the observed anti-inflammatory effects, these findings are consistent with the hypothesis that Sig-1R activation mediates the DMT-induced phenotypic shift in microglia.

To further investigate the mechanism underlying the inhibitory effects of DMT on microglial activation and to test the alternative hypothesis that DMT exerts its anti-inflammatory effects through 5-HT receptor activation (Krabbe et al., 2012; de Deus et al., 2025), we next pharmacologically inhibited Sig-1Rs and 5-HT receptors in the presence of DMT in cultured microglia and acute brain slices. Unexpectedly, the DMT-induced shift of microglia toward a homeostatic morphological phenotype required activation of both Sig-1Rs and 5-HT receptors (Fig. 6, Fig. 9), whereas its effects on phagocytic activity were largely independent of either signaling pathway (Fig. 7). Morphological activation is primarily driven by cytoskeletal reorganization (Nolte et al., 1996), while phagocytosis depends on specialized recognition and engulfment machinery (Paolicelli et al., 2022). The differential effects of receptor antagonism observed in the present study support the emerging concept that these processes constitute related but mechanistically distinct programs of microglial activation, rather than inseparable manifestations of a single activated state.

Our findings are also consistent with previous studies. In a stress-induced hypertension rodent model complemented by primary or BV2 microglial cultures, the synthetic Sig-1R agonist SKF10047 attenuated pro-inflammatory microglial polarization, supporting a role for Sig-1R activation in regulating microglial inflammatory responses (Ooi et al., 2021). Similarly, serotonin reuptake inhibitors, which increase extracellular serotonin concentrations and thereby enhance serotonergic signaling, have also been shown to exert anti-inflammatory effects on microglia (Liu et al., 2011; Tynan et al., 2012). Together, these findings support the possibility that DMT engages both Sig-1Rs and 5-HT receptors to promote a more homeostatic microglial morphology. In contrast, experimental evidence for a role of Sig-1Rs in regulating microglial phagocytosis is scarce, and the available data on serotonergic regulation of microglial phagocytosis are inconsistent. For example, serotonin reuptake inhibitors, which suppress the pro-inflammatory polarization of microglia (Liu et al., 2011; Tynan et al., 2012), have been reported to enhance microglial phagocytosis (Park et al., 2021). Conversely, an earlier study demonstrated that serotonin itself reduced phagocytic activity (Krabbe et al., 2012). In the present study, the suppressive effect of DMT on phagocytosis was not reversed by inhibition of either Sig-1Rs or 5-HT receptors, suggesting that the mechanisms regulating phagocytosis differ, at least in part, from those underlying DMT-induced morphological reprogramming of microglia.

The brain slice experiments extend our findings beyond the direct effects of DMT on microglia by demonstrating that DMT also protects against SDs and ischemic neuronal injury (Fig. 8), in agreement with previous studies demonstrating the neuroprotective effects of DMT in experimental ischemic stroke (Nardai et al., 2020; Szabó et al., 2021; László et al., 2025). Pharmacological receptor profiling further revealed that this tissue-level neuroprotection depends on serotonergic signaling, at least in the present experimental model (Fig. 8). These novel observations complement and extend previous evidence implicating Sig-1Rs in DMT-mediated neuroprotection. Specifically, co-administration of the Sig-1R antagonist BD1063 abolished the infarct-limiting effect of DMT (Nardai et al., 2020), and the Sig-1R antagonist NE-100 mitigated the inhibitory effect of DMT on SDs (Szabó et al., 2021). By contrast, our findings indicate that serotonergic signaling also contributes to DMT-mediated neuroprotection, highlighting a previously underappreciated receptor mechanism. Taken together, the available evidence suggests that the receptor mechanisms underlying the consistently observed neuroprotective effects of DMT in experimental ischemic brain injury are context dependent, with the relative contributions of Sig-1R and serotonergic signaling varying according to the experimental model, cellular target, and biological endpoint examined.

Finally, given the rapid clinical translation of DMT (Stojanović et al., 2026), including its development as a potential therapy for ischemic stroke (László et al., 2025; van der Heijden et al., 2025), understanding its mechanisms of action has become increasingly important. Our study provides new mechanistic insights into the cellular and receptor-mediated actions of DMT in the injured brain that may help guide its future clinical development.

## LIMITATIONS AND FUTURE DIRECTIONS

There are several limitations inherent to the technical approaches used in this study. Primary microglial cultures are derived from the neonatal neocortex and therefore may not fully recapitulate the mature phenotype of adult microglia. Nevertheless, they remain a widely used experimental model for investigating microglial biology and pharmacological modulation (Pesti et al., 2024). In addition, the proteomic analyses performed in this study quantified changes in protein abundance rather than enzymatic activity. Consequently, future studies employing enzyme activity assays will be required to determine the functional significance of the identified proteins. Finally, receptor mechanisms were investigated using pharmacological antagonists. Because asenapine antagonizes multiple serotonin receptor subtypes with high affinity, the present experiments cannot identify the individual 5-HT receptor subtype(s) responsible for the anti-inflammatory and neuroprotective effects of DMT. Further studies using highly selective receptor antagonists or genetic receptor knockout approaches will therefore be required to define the precise serotonergic mechanisms involved.

The present study focused exclusively on acute injury and acute DMT treatment. Although this approach is well suited to mechanistic investigations, it does not address the consequences of prolonged DMT administration or the long-term effects on microglial function and ischemic brain injury. Future *in vivo* studies employing chronic DMT treatment paradigms, such as those recently introduced (László et al., 2025), will therefore be important to establish the translational relevance of these findings. Finally, DMT is rapidly metabolized by monoamine oxidases, resulting in a short biological half-life (Schimmelpfennig and Jankowiak-Siuda, 2025). Consequently, co-administration of monoamine oxidase inhibitors may prolong DMT exposure and potentiate its therapeutic effects (Egger et al., 2024), an aspect that also warrants further investigation.

## DATA AVAILABILITY

The mass spectrometry proteomics data generated in this study have been deposited to the ProteomeXchange Consortium via the PRIDE partner repository with the dataset identifier PXD082149. The dataset will remain private during peer review and will be made publicly available upon publication. The scripts used to analyze the data and generate the figures are available from the corresponding author upon reasonable request.

## AUTHOR CONTRIBUTIONS (CRediT STATEMENT)

I. P., Á.B and R.F.: Investigation, Data curation, Formal Analysis, Visualization, Writing – review & editing; Z.D.: Investigation, Data curation, Formal Analysis, Visualization, Funding acquisition, Writing – review & editing; S.D.: Investigation, Formal Analysis, Visualization, Funding acquisition, Writing – review & editing; Z.P.: Data curation, Formal Analysis, Visualization, Writing – review & editing; T.P.: Funding acquisition, Writing – review & editing; É.H-G.: Investigation, Methodology, Writing – review & editing; F.B.: Supervision, Writing – review & editing; Á.M.: Methodology, Supervision, Writing – review & editing; K.K., S.P. and K.V.: Investigation, Data curation, Visualization, Writing – review & editing; N.V.C.: Conceptualization, Investigation, Writing – review & editing; E.F.: Conceptualization, Data curation, Formal Analysis, Visualization, Writing – original draft, Funding acquisition.

